# The FliI ATPase couples ATP hydrolysis to substrate switching in bacterial flagellar type-III secretion

**DOI:** 10.1101/2025.08.04.668468

**Authors:** Rosa Einenkel, Mario Delgadillo-Guevara, Lasse Hallenga, Christian Goosmann, Marc Erhardt

## Abstract

Bacterial flagella are assembled by a specialized type III secretion system that exports structural subunits in a defined order. While the ATPase FliI is known to couple ATP hydrolysis to substrate translocation, its role in the transition between early and late secretion stages has remained unclear. Here, we systematically analyzed *Salmonella enterica* strains with defined FliI point mutations and found that FliI activity is dispensable for early substrate export and hook-basal body formation but is important for triggering the substrate specificity switch and promoting late substrate export. Mutant strains showed delayed gene expression from class 3 promoters, prolonged early secretion, and impaired flagellar filament assembly, despite normal FliI localization and oligomerization. These findings support the involvement of FliI in controlling the temporal dynamics of flagellar assembly. We propose that FliI contributes to substrate switching, ensuring robust and orderly fT3SS function. This study highlights the multifaceted role of the fT3SS ATPase in optimizing the efficiency and robustness of flagellum assembly.

**Significance:** The ordered export of substrates by bacterial type III secretion systems is essential for the assembly of complex surface structures such as the flagellum, yet the mechanisms that control the timing of substrate switching remain poorly understood. The bacterial flagellar ATPase FliI is best known for being involved in energizing the flagellar type III secretion system. Here, we show that FliI contributes to the transition from early to late substrate export during flagellar biogenesis. Using targeted FliI mutants in *Salmonella*, we show that ATPase activity is dispensable for early export but important for proper substrate switching and late-stage flagellar assembly. These findings highlight the multifaceted roles of FliI during flagellar assembly beyond activating the flagellar type III secretion system.

## Introduction

Bacterial flagella are complex nanomachines that play a crucial role in motility, enabling bacteria to navigate through their environment (1). This motility is essential for many bacterial processes, including colonization and adhesion to host cells (2). The flagellum consists of three major structural components: the basal body, the hook and the filament (**Fig. 1A**). The basal body (BB) anchors the flagellum to the bacterial cell envelope and serves as the motor apparatus, generating the rotational force that drives motility (3, 4). It is composed of several ring-like structures, including the peptidoglycan-spanning P-ring the outer membrane L-ring and the inner membrane MS-ring, which connects to the rotor (3, 4). The rotor, or C-ring links the flagella mediated motility to the process of chemotaxis and acts as the rotary switch complex (5–7). The stator units generate the rotational force that drives the rotation of the C-ring and hence the flagellar filament (8). The proximal rod and the distal rod form the secretion channel through the cell envelope. The hook acts as a flexible joint between the BB and the filament (9), with its length precisely regulated (10, 11). The hook-filament junction connects the hook to the filament, which consists of tens of thousands of subunits of a single protein, the flagellin (either FliC or FljB in *S. enterica*) and acts as a mechanical buffer zone to smoothly connect the flexible hook and the rigid filament (12). The filament’s distal end is capped by a pentamer of FliD, a specialized structure, that facilitates the assembly of flagellin subunits into a functional filament (12, 13).

**Fig. 1.**
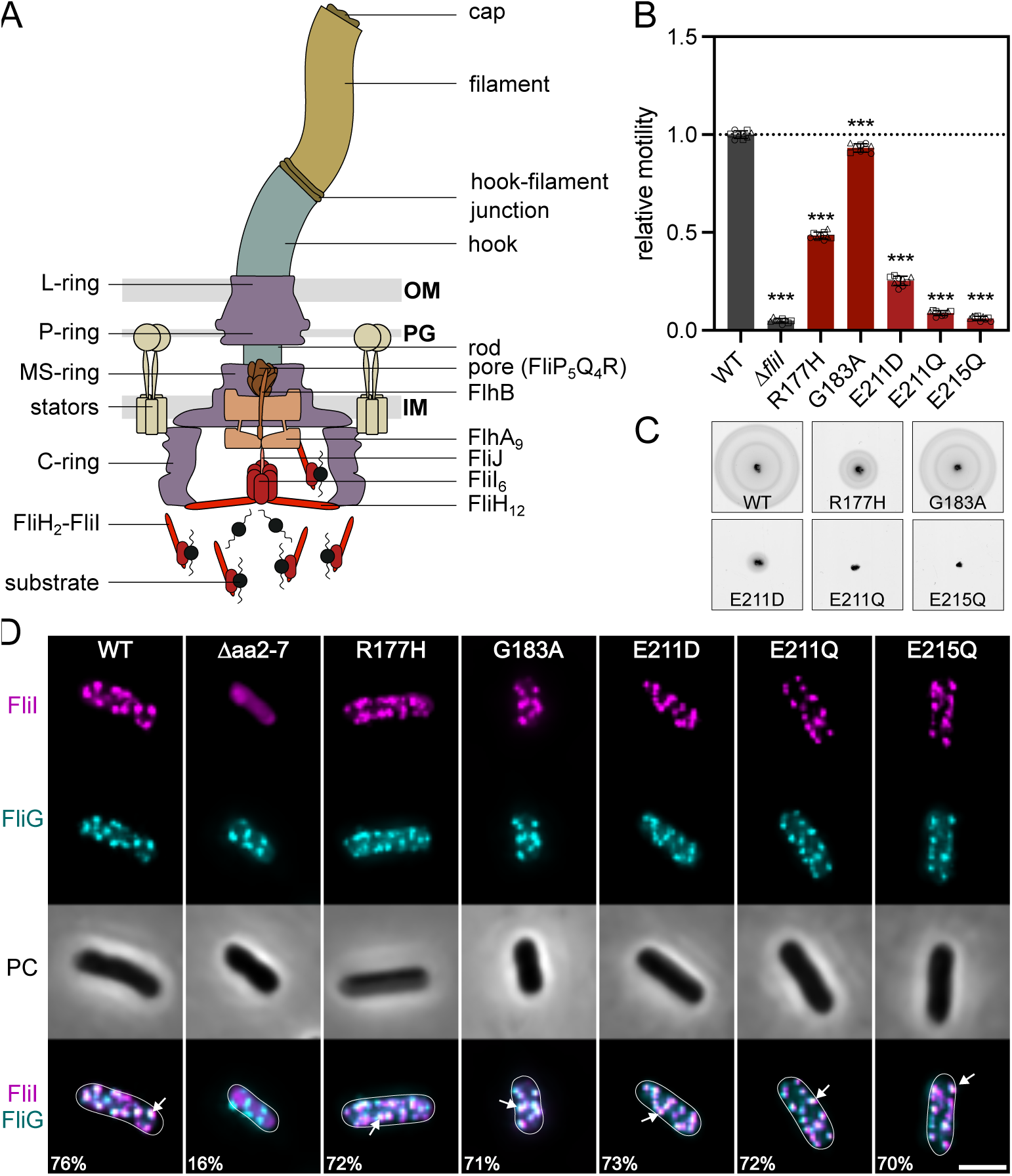
FliI is required for flagellar motility and efficient assembly of the flagellar export apparatus. (A) Schematic of a gram-negative bacterial flagellum and the flagellar type III secretion system (fT3SS). The flagellar motor spans the inner membrane (IM), periplasmic space including the peptidoglycan (PG), and outer membrane (OM), and assembles an external filament via a central secretion channel. The cytoplasmic ATPase complex (FliHIJ) delivers substrates and activates the export gate through ATP hydrolysis. (B) Relative motility of wildtype (WT) and *fliI* mutants analyzed using soft-agar motility plates, quantified after 4-5 h incubation at 37 °C. Diameters of the motility halos were measured using Fiji and normalized to the WT. Bar graphs represent the mean of at least 3 biological replicates with standard deviation error bars. Replicates shown as individual data points. ***p < 0.001 (ordinary one-way ANOVA with Dunnett’s multiple comparison test vs. WT). (C) Representative swimming halos of the WT and *fliI* point mutants. (D) Representative microscopy images of WT and *fliI* mutants. FliI-HaloTag labeled with TMR-ligand (magenta), basal bodies visualized with mNeonGreen-FliG (cyan); PC (PhaseContrast). Percentage of FliI-FliG foci pairs below the colocalization threshold (2 px) is indicated for each strain. White arrows show examples of colocalized foci. Scale-bar: 2 µm.

The secretion of axial proteins is tightly regulated and mediated by the flagellar type III secretion system (fT3SS), which is responsible for the translocation of most extra-cytoplasmic flagellar components from the bacterial cytoplasm to the site of assembly, a process that is mainly powered by the proton motive force (14). The fT3SS as part of the BB consists of the membrane embedded export gate (FlhA, FlhB) and secretion pore (FliP, FliQ and FliR), as well as a cytoplasmic ATPase complex (FliH, FliI and FliJ) (15). FliI, a Walker-type AAA^+^-ATPase with two highly conserved motifs for nucleotide binding (Walker A) and ATP hydrolysis (Walker B), forms a hexameric ring structure at the base of the flagellum with FliJ as the central stalk, anchored to the basal body by 12 FliH molecules through interactions with the motor protein FliN (16–21). The ATPase complex is thought to couple ATP hydrolysis to the activation of the export gate through interactions of FliJ with the export gate protein FlhA to ensure efficient and robust protein export (22). While a previous study had proposed a potential interaction between the second messenger c-di-GMP and the fT3SS ATPase FliI (23), the exact role of this interaction remains unclear. Furthermore, heterotrimers of 2 FliH and 1 FliI are thought to play an important role in the dynamic delivery of substrates to the export gate, ensuring the timely assembly of the flagellar components (24). Based on the known functions of the evolutionarily related ATPase InvC from the virulence-associated T3SS, as well as insights from mathematical modeling, FliI has been proposed to play roles in substrate unfolding and in dissociating substrates from their cognate chaperones (25, 26). However, although extensively studied, the exact molecular mechanisms of the fT3SS ATPase complex remain incompletely understood (27).

Flagellum biogenesis and flagellar gene expression are tightly regulated through a hierarchical system to ensure proper assembly and efficient energy utilization. This process involves three distinct classes of promoters, starting from the class 1 promoter that drives the expression of the master regulator FlhD_4_C_2_. FlhD_4_C_2_ activates expression from class 2 promoters, that encode structural components of the hook-basal body (HBB) as well as regulatory proteins like the flagellar-specific transcription factor σ^28^ and its anti-σ factor FlgM. FlgM coordinates the temporal progression of flagellar gene expression by sequestering σ^28^, thereby inhibiting transcription from class 3 promoters until hook assembly is completed. Hook completion acts as a checkpoint that signals the export apparatus to switch substrate specificity from early substrates (rod and hook components) to late substrates (filament components). This transition, mediated by FlhB and FliK, ensures proper hook length, which is essential for optimal motility and flagellar bundle formation (10, 11, 26). Late substrates require chaperones for their efficient export and to prevent premature interactions and degradation. Upon the substrate specificity switch, FlgM is recognized as a late substrate and secreted via the fT3SS, freeing σ^28^ to finally initiate transcription from class 3 promoters, thereby elegantly connecting proper flagellum assembly with gene regulation (25, 28, 29). Genes expressed from class 3 promoters encode, among others, the flagellins, the stator units and key components of the chemotaxis system (30).

In this study, we dissected the role of the flagellar ATPase FliI by generating targeted point mutations in its ATPase domain and adjacent regions implicated in secretion control. Through a combination of mutagenesis and functional assays, we identified mutants that exhibited selective defects in the secretion of late substrates and in the substrate specificity switch, while early substrate export and hook assembly remained unaffected. These results demonstrate that FliI is dispensable for early stages of flagellar assembly but important for triggering the transition to late-stage export. Our findings support a model in which FliI not only powers secretion via ATP hydrolysis but also actively supports the substrate switching mechanism, thereby ensuring the temporal fidelity and efficiency of flagellar biogenesis.

## Results

The Walker-type AAA+ ATPase FliI from *S. enterica* plays a crucial role in flagellar assembly by ensuring a robust export of substrates through the fT3SS. To investigate the molecular function of FliI, we generated point mutations at residues implicated in its catalytic activity and putative regulatory sites (23, 31). All mutations were introduced into the *S. enterica* genome to maintain native expression conditions. Specifically, we created mutations at residues located at the interface between FliI monomers (R177H and E215Q), and in the nucleotide-binding Walker A motif within the central cavity of the FliI hexamer (G183A). Additionally, E215 was previously shown to be involved in the catalytic activity of FliI (19). We included two previously characterized Walker B mutants (E211D and E211Q), known to drastically reduce ATPase activity - by 100-fold and completely, respectively - resulting in impaired motility and substrate secretion (31).

The effect of *fliI* point mutations on swimming motility was assessed using a soft-agar motility assay (**Fig. 1B-C**). As anticipated, the *fliI* deletion mutant (Δ*fliI*) was non-motile, consistent with previous reports (14). The R177H mutant displayed 48% motility, while the E215Q mutant was non-motile. The Walker A motif mutant G183A exhibited a slight reduction in motility to 93% of the WT level. The Walker B motif mutants E211D and E211Q showed 25% motility or loss of motility, respectively. These phenotypes suggest defects in flagellar assembly, prompting further investigation into how these specific mutations impair motility.

To investigate whether the phenotypes observed in the FliI mutants were caused by subcellular mislocalization of the ATPase, we performed *in vivo* fluorescence microscopy. In order to observe FliI localization, we constructed strains expressing FliI with a C-terminal HaloTag fusion. To assess colocalization of the ATPase with the basal body, the strains also expressed the C-ring component FliG with an N-terminal mNeonGreen fusion. Flagellar biosynthesis was synchronized by replacing the native promoter of the *flhDC* master regulator operon with an anhydrotetracycline (AnTc) inducible P*_tetA_* promoter (32). FliI-HaloTag was labeled with the cell-permeant ligand tetramethylrhodamine (TMR), and fluorescence microscopy was performed on live cells (**Fig. 1D**). We then quantified the spatial colocalization between foci detected in the respective fluorescence channels by calculating pairwise distances between detected spots across individual cells. Spots were considered colocalized if their distance was below a predefined threshold (2 px or ∼144 nm). The fraction of colocalized foci pairs was calculated as the number of colocalized pairs divided by the total number of detected pairs within a maximal distance (30 px or ∼2µm).

In WT cells, FliI strongly colocalized with FliG at the basal body, forming distinct overlapping foci with 76% of foci pairs below the colocalization threshold (**Fig. 1D**). As a negative control, the FliI Δaa2-7 mutant, which has been previously shown to be deficient in hexamer formation (33) and interaction with FliH (34), displayed a diffuse cytoplasmic localization, with no distinct FliI foci visible by microscopy. Due to the nature of the analysis pipeline, which detects local intensity maxima, foci pairs were still found, however only 16% were considered colocalized. This confirms that FliI Δaa2-7 is unable to localize to the basal body, likely due to its inability to properly form hexamers and disrupted interaction with FliH. The FliI point mutants (R177H, G183A, E211D, E211Q, E215Q) localized to the basal body similarly to the WT, as indicated by the presence of discrete FliI foci (Figure 4D). The percentage of colocalized foci pairs was 72% for R177H, 71% for G183A, 73% for E211D, 72% for E211Q, and 70% for E215Q. These results suggest that the FliI point mutations do not substantially impair the subcellular targeting of FliI. Additionally, we confirmed that proper localization of FliI depends on FliH and FliN but not on FliJ or FlhA, as previously reported (**Fig. S1A**) (21).

To assess whether the FliI point mutants affect hexamer formation, we performed *in vivo* site-specific photo-crosslinking of FliI variants containing a photoactivatable pBpa residue (FliI*) at the FliI-FliI interface (**Fig. S1B**). UV-dependent formation of higher molecular weight FliI adducts was observed for WT FliI* and all its point mutants studied here, indicating that these variants retain the ability to oligomerize (**Fig. S1C**). In contrast, FliI* Δaa2-7 exhibited only a faint high molecular weight band, suggesting premature oligomerization in the cytoplasm, likely due to the high-level plasmid expression of FliI*. However, the reduced level of cross-links compared to the WT confirms that hexamer formation in this mutant is defective but may still occur at low levels under the experimental conditions. Taken together, these results demonstrate that while the mutations in FliI impair motility to varying degrees, they do not substantially alter its subcellular localization or ability to form oligomers. Thus, the observed motility defects in the FliI point mutants may instead reflect disruptions in the secretion activity of the fT3SS.

### FliI facilitates secretion of late flagellar substrates

The fT3SS mediates the assembly of the axial structures of the flagellum through two distinct secretion modes: early and late. Upon hook completion, the fT3SS undergoes conformational changes, allowing recognition and export of late substrates (11, 35). To determine whether the motility defects of the FliI point mutants result from impaired secretion, we systematically analyzed early and late substrate export in these strains.

First, we confirmed FliI localization in strains deleted for components that lock the fT3SS in either the early (Δ*flgBC* (proximal rod), Δ*flgE* (hook)) or late (Δ*flgKL* (hook-filament junction)) stages of flagellar assembly (**Fig. S1A**). Fluorescence microscopy revealed that FliI colocalized with the basal body in both secretion states, indicating its presence at the basal body throughout the flagellar assembly process. To analyze the secretion efficiency of early substrates in FliI point mutants, we employed a hook protein / β-lactamase (FlgE-Bla) reporter system (**Fig. 2A**) (36, 37). Deletion of the rod components FlgBC leads to secretion of flagellar substrates into the periplasm (38), resulting in the ability to resist ampicillin in correlation to the quantity of secreted FlgE-Bla. Therefore, we analyzed the ability of the FliI mutants to grow in the presence of increasing ampicillin concentrations and determined the concentration of ampicillin leading to inhibition of bacterial growth to half of its maximum (IC_50_ [Amp], **Fig. S2A**) and then normalized it to WT levels (**Fig. 2B**). A mutant lacking the MS-ring and therefore the entire fT3SS (Δ*fliF*), was used as a negative control, representing the basal level of resistance against ampicillin (**Fig. S2A**). Deletion of *fliI* resulted is a severe reduction of early substrate secretion to 2% of WT levels, similar to the Δ*fliF* control strain (1%), consistent with previous observations (14).

**Fig. 2.**
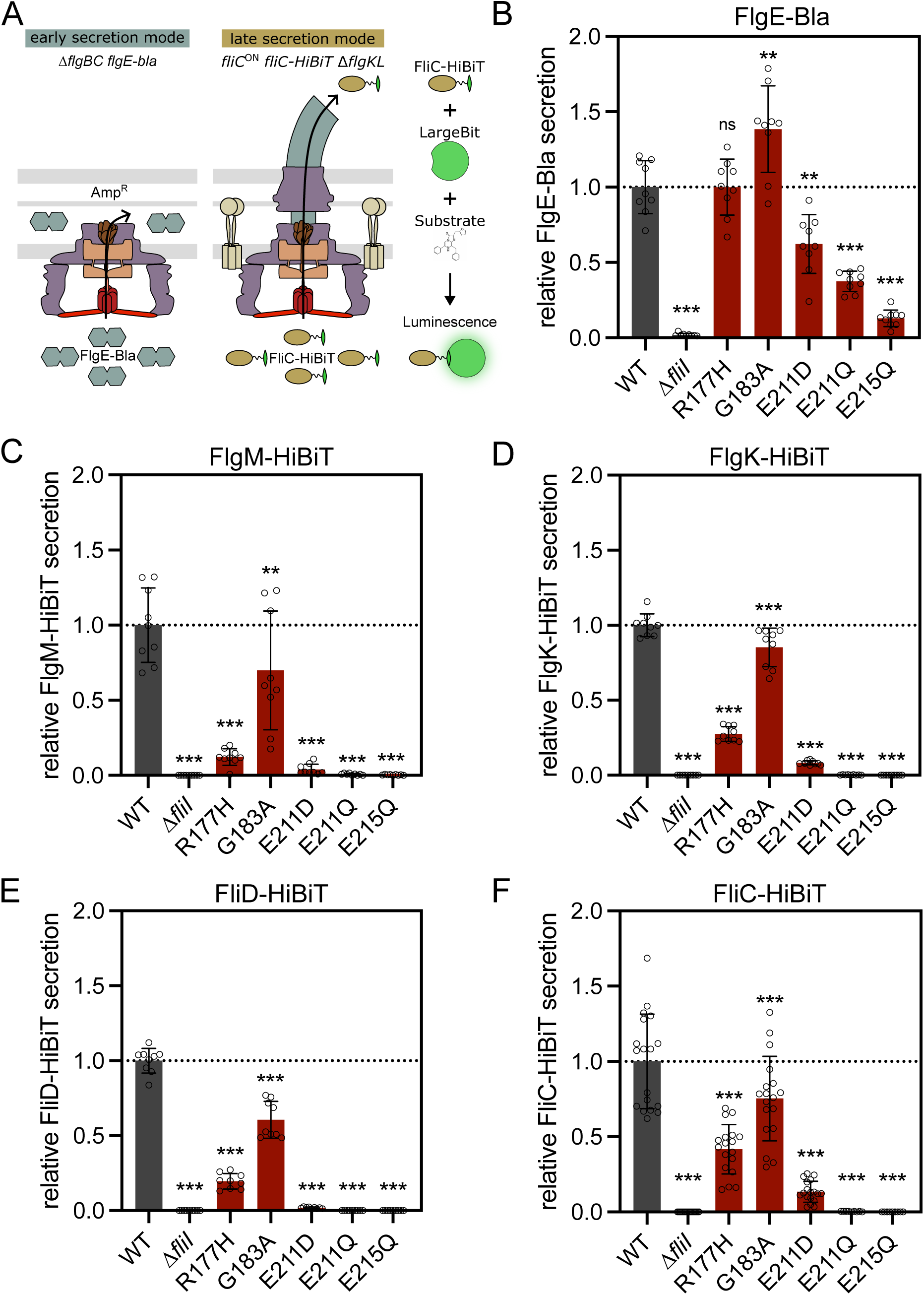
FliI facilitates secretion of late substrates. (A) Schematic of experimental set-ups to quantify secretion of early (FlgE-Bla) and late substrates (example for FliC-HiBiT). Left: early substrate secretion was measured in strains lacking proximal rod components, allowing secretion of FlgE-Bla into the periplasm, where it confers ampicillin-resistance (Amp^R^), proportional to the amount of secreted FlgE-Bla. Right: late substrate secretion was quantified using the split Luciferase NanoBit. Strains were locked in FliC production (*fliC*^ON^) and lacked the hook-filament junction to prevent filament assembly (Δ*flgKL*). Cells expressed a translational fusion of FliC to HiBiT. Addition of LargeBit and substrate to the culture supernatant enabled luminescence detection proportional to secreted FliC-HiBiT levels. (B) Quantification of early substrate (FlgE-Bla) secretion. Relative ampicillin resistance normalized to WT. Bars represent means ± SD from at least 3 independent experiments; **p < 0.01, ***p < 0.001; ns, not significant (ordinary one-way ANOVA with Dunnett’s multiple comparison test vs. WT). (C-F) Quantification of late substrate secretion, normalized to the WT. Bars represent means ± SD from at least 3 independent experiments; ***p < 0.001 (ordinary one-way ANOVA with Dunnett’s multiple comparison test vs. WT).

Among the point mutants, G183A exhibited the highest secretion efficiency, with an IC_50_ of 356 µg/mL corresponding to 138% of WT levels. The R177H mutant displayed secretion comparable to WT (IC_50_ of 257 µg/mL compared to 256 µg/mL for WT), despite reduced significantly motility. Thus, both R177H and G183A were proficient in early substrate secretion, despite motility defects. In contrast, the ATPase activity mutants showed impaired secretion. E215Q exhibited the most severe defect, with only 13% of WT levels (32 µg/mL), though still 11-fold higher than the Δ*fliF* control. The Walker B motif mutants E211D and E211Q retained partial secretion activity (62% and 37% of WT, respectively). Importantly, this still represents a significant 52- and 31-fold increase of the IC_50_ of the ATPase activity mutants E211D (IC_50_ = 157 µg/mL) and E211Q (IC_50_ = 95 µg/mL) compared to the Δ*fliF* control strain, respectively (**Fig. S2A**).

For quantification of late substrate secretion, we next employed the Nano-Glo extracellular detection system (Promega) and the split luciferase NanoBit (**Fig. 2A**) (39, 40). Strains were engineered to express late flagellar substrates translationally fused to the small luciferase subunit HiBiT (41). Secreted HiBiT-tagged proteins accumulated in the supernatant and were detected via luminescence (**Fig. 2C-F**).

All mutants with impaired early secretion (Δ*fliI*, E211D, E211Q, and E215Q) also exhibited strong defects in the secretion of FlgM-HiBiT (**Fig. 2C**). Secretion levels were reduced to 0.1% (Δ*fliI*), 0.4% (E215Q), 0.7% (E211Q) and 7% (E211D) of WT levels. The R177H mutant also showed a significant reduction (12.3%), while G183A retained 69.9% of WT levels. FlgK-HiBiT secretion (**Fig. 2D**) was also reduced across all mutants. The strongest defects were observed in *ΔfliI*, E215Q, and E211Q, each exhibiting less than 0.3% of WT levels. E211D retained 8.2%, while R177H and G183A secreted 27.5% and 85.2% of WT levels, respectively.

A similar pattern was observed for FliD-HiBiT (**Fig. 2E**), with secretion entirely abolished in *ΔfliI*, E215Q, and E211Q (0.0%). E211D retained minimal export (1.8%), and secretion was strongly to moderately reduced in R177H (19.6%) and G183A (60.5%).

E211D retained partial FliC-HiBiT (**Fig. 2F**) secretion (13.5% of WT), while the Δ*fliI* and E215Q mutants showed only 0.1%, and E211Q 0.3% of WT levels. Secretion of FliC-HiBiT was moderately reduced to 75.4% in G183A and more strongly in R177H (41.7%), consistent with their motility phenotypes. Western blot analysis of cellular and secreted fractions (**Fig. S2B–E**) confirmed the luminescence data. Additionally, reduced cellular levels of FliC-HiBiT in most mutants suggested a delayed substrate specificity switch, likely contributing to the impaired secretion phenotypes (**Fig. S2E**).

Secretion of all tested late substrates was consistently reduced across the *fliI* mutants, suggesting a general requirement for FliI in late substrate export. Together, these results are consistent with previous reports that early substrates can be secreted with minimal ATP hydrolysis (31). The increased early substrate secretion in the G183A mutant, and the unaffected secretion in the R177H mutant, indicate that FliI activity is likely more critical for late substrate export than for early stages of assembly. This is further supported by the observation that all ATPase activity mutants (E211D, E211Q and E215Q) retained partial early secretion, despite strongly impaired ATPase activity. Together, these findings suggest that FliI may promote late substrate export secretion either by enhancing transport efficiency or by contributing to the substrate specificity switch.

### FliI contributes to the substrate specificity switch

The observed decrease of late substrate secretion in the *fliI* mutant strains could have resulted from either a defect in secretion efficiency of late substrates or from a defect or delay in the substrate specificity switch. Before hook completion, the late substrate and anti–ο^28^ factor FlgM inhibits the transcription of genes from class 3 promoters by complexing the flagellar specific ο^28^ (42). The fT3SS undergoes a substrate specificity switch upon complete formation of the HBB, which results in the secretion of FlgM, thereby allowing formation of the ο^28^-RNA-Polymerase holoenzyme and transcription from class 3 promoters (43). Inefficient late substrate secretion would therefore not only result in lower FliC secretion levels but simultaneously result in a lower abundance of late substrates due to prolonged sequestration of ο^28^ by FlgM.

To address possible delays in the switch mechanism, gene expression from class 3 promoters was monitored using strains with an inducible *flhDC* master operon for synchronized flagella formation and a plasmid harboring the reporter genes *luxCDABE* under control of the class 3 promoter P*_motA_* (44). We found for the WT a half-maximal induction time of 93 ± 3 min of the P*_motA_* promoter after *flhDC* induction (**Fig. 3A**, dotted line). Further, the relative class 3 gene expression at T_90_ and T_360_ was determined (**Fig. S2A**). The previously described slow hook polymerization mutant *flgE*_T149N_ was utilized as a control (45, 46). Since the hook polymerization in this mutant takes much longer, the substrate specificity switch and therefore activation of transcription from class 3 promoters is delayed.

**Fig. 3.**
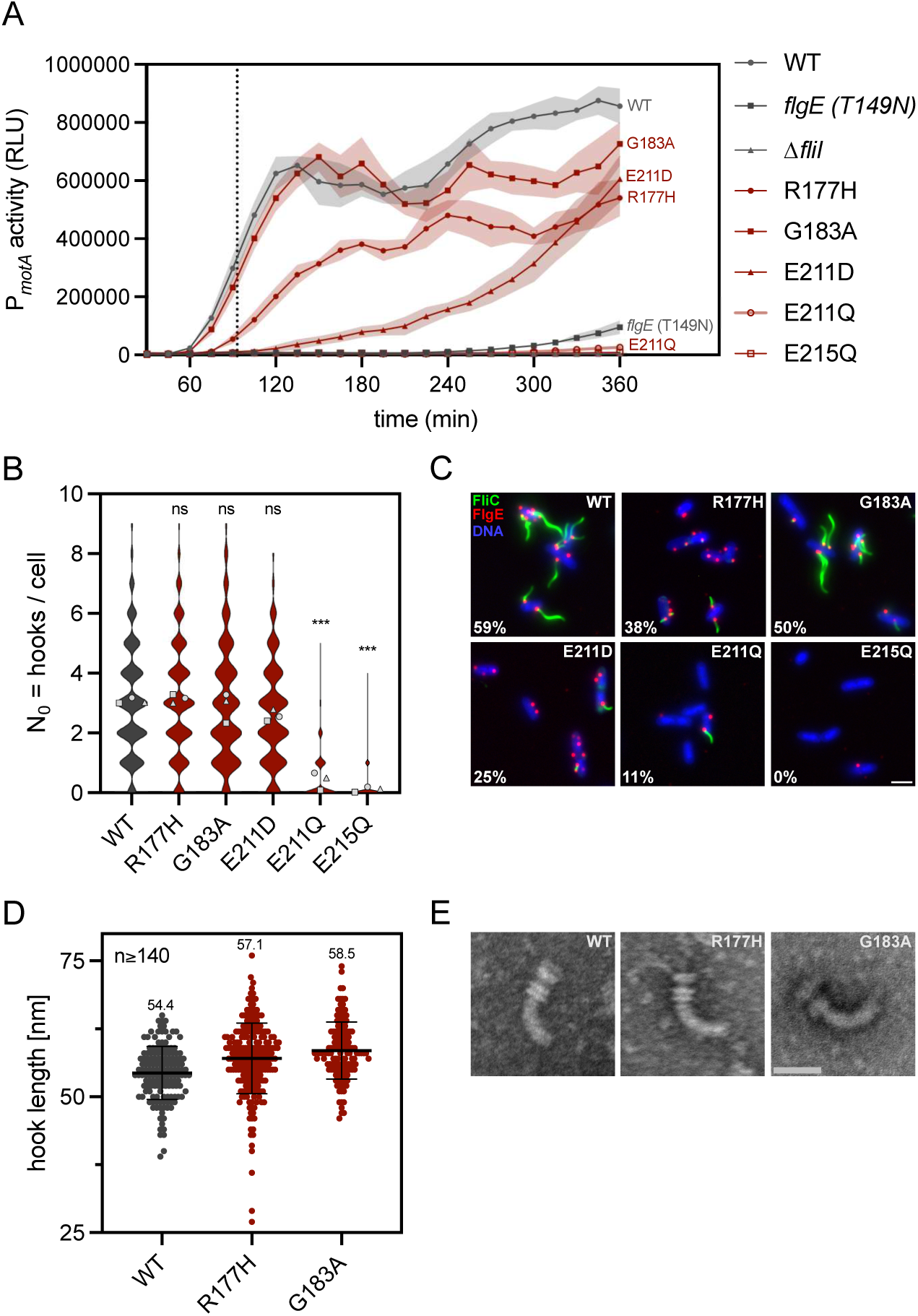
FliI contributes to the substrate specificity switch. (A) Time course of P*_motA_* activity (class III gene expression). Luciferase reporter activity was monitored in strains with synchronized flagella synthesis. Shown are the mean RLU (lines and symbols) ± SD (transparent lines) measured in 6 biological replicates. Dashed line indicates half-maximal induction time for P*_motA_* in the wildtype (WT). (B) Quantification of hook number per cell determined by fluorescence microscopy. Violin plots represent distribution of the data from 3 biological replicates; symbols represent the mean of the biological replicates. ***p < 0.001; ns, not significant (ordinary one-way ANOVA on the means with Dunnett’s multiple comparison test vs. WT). (C) Representative fluorescence microscopy images of WT and *fliI* point mutants. Filaments (FliC T237C) labeled with Star Green Maleimide (green), hooks (FlgE_3×HA_) immunostained with α-HA-Alexa555 (red) and DNA counterstained with DAPI. Percentage of hooks with an attached filament indicated in white. Scale-bar: 2 µm. (D) Hook length measurements from purified hook basal bodies. Symbols represent individual measurements, horizontal black line indicates mean ± SD error bars. (E) Representative negative-stain Transmission Electron Microscopy images of purified hook basal bodies. Scale bar = 50 nm.

**Fig. 4.**
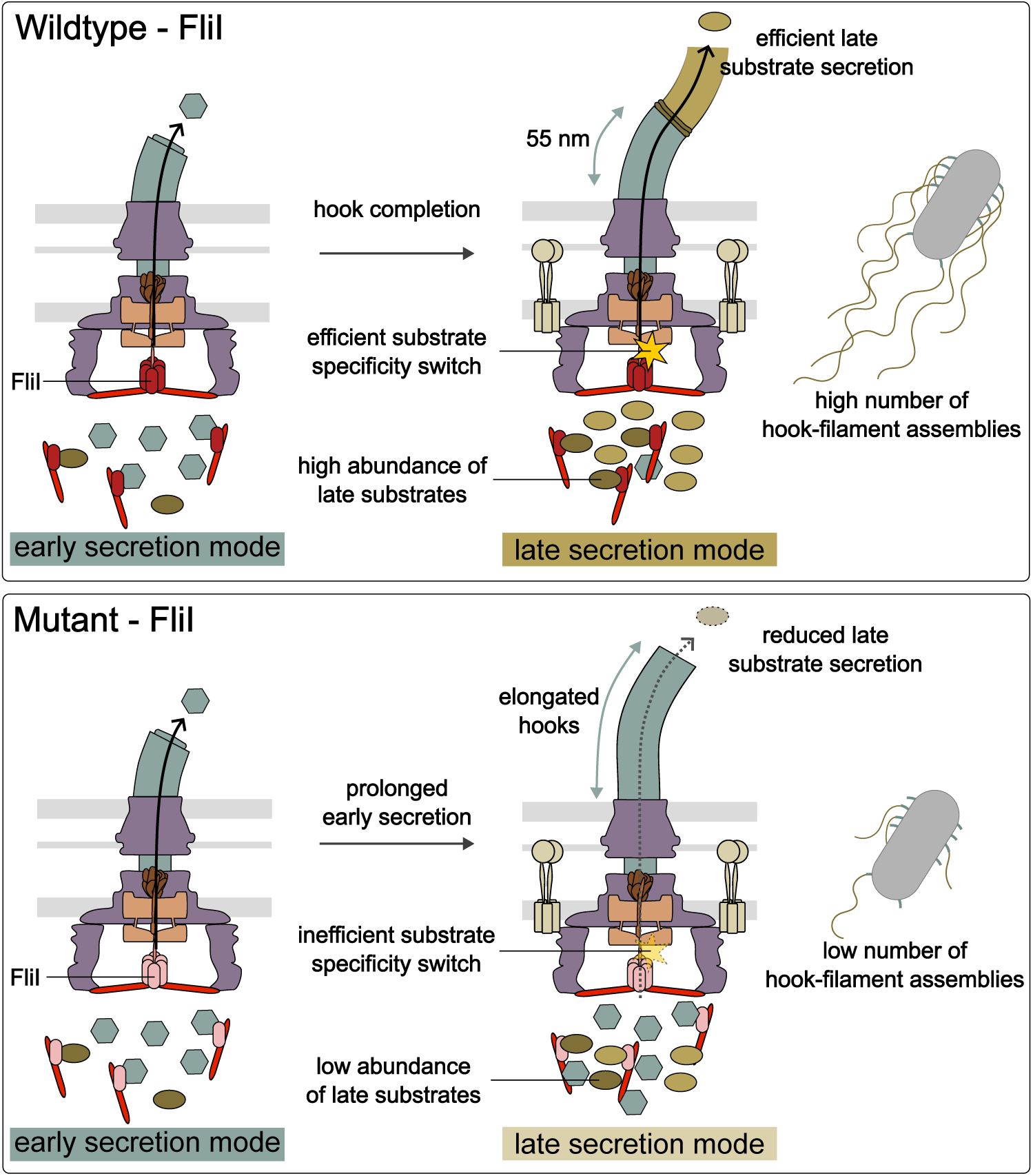
Schematic model of FliI function in wildtype and mutant strains. In wildtype cells (top), FliI facilitates the transition from early to late secretion upon hook completion. This enables efficient FlgM secretion, timely induction of class 3 gene expression, and robust export of late substrates, resulting in many hooks with attached filaments. In FliI mutants (bottom), prolonged early secretion, delayed FlgM export, impaired class 3 gene expression, and inefficient late substrate export are observed. Consequently, hooks are elongated, and fewer filaments are assembled.

The G183A mutant displayed the mildest defect in the capability to perform the substrate specificity switch with a relative class 3 gene expression of 78% and 85% at T_90_ and T_360_, respectively. While P*_motA_* showed only 18% of the WT activity at T_90_ in the R177H mutant, the relative promoter activity reached 63% at T_360_. The ATPase activity mutant E211D showed a steady increase of relative class 3 gene expression over time, starting from 3% at T_90_ to 71% at T_360_. The E211Q mutant displayed a relative class 3 gene expression of 3% at T_90_ and T_360_. In both Δ*fliI* and E215Q mutants the relative class 3 gene expression stayed approximately the same over the course of the experiment, with 2% (T_90_) and 1% (T_360_) relative expression observed. The *flgE*_T149N_ mutant showed a delayed substrate specificity switch, as expected, resulting in relative class 3 gene expression of 2% at T_90_ and 11% at T_360_.

A possible reason for the observed reduction of class 3 gene expression in the mutant strains could be a reduced number of hooks. A slower hook polymerization, similar to the *flgE*_T149N_ control strain, and hence a reduced number of completed hooks would delay the substrate specificity switch. Only upon completion of the HBB, the substrate specificity switch occurs and the anti-σ28 factor FlgM will be recognized as a substrate of the fT3SS. Accordingly, a delayed and/or decreased HBB formation frequency, resulting in FlgM remaining inside the cell, would also explain the observed delay in class 3 gene expression.

To address this possible reason for the observed delayed substrate specificity switch in the *fliI* mutants, we determined the number of hooks per cell (**Fig. 3B**). This was especially interesting for the R177H and G183A mutant, since both mutants did not display a reduction of early substrate secretion but showed reduced class 3 gene expression levels (**Fig. 2C**, **Fig. 3A**). Furthermore, we determined the number of HBBs with an attached filament (**Fig. 3C**, **Fig. S4**). We hypothesized, if the substrate specificity switch was intact, that the ratio of attached filaments to the HBBs would be similar to the WT. On the other hand, an increased number of HBBs without attached filaments would suggest an impairment in the substrate specificity switch. To determine the number of hooks and attached filaments per cell, we simultaneously visualized the hook and filament using strains expressing *flgE*_3×HA_ for immunostaining of the hook and *fliC*_T237C_ for maleimide staining of the filament (**Fig. 3C**). In agreement to the early substrate secretion levels, both R177H and G183A mutants did not show a difference in the amount of assembled HBBs compared to the WT. The ATPase activity mutant E211D displayed a slight, but non-significant reduction of hooks per cell. In contrast, E211Q displayed a significantly reduced number of hooks with a maximum of five hooks per cell, compared to nine in the WT. The number of cells without hooks just increased slightly from 8% (WT) to 10% in E211D, whereas 55% of the E211Q mutant did not assemble any hooks. The number of hooks per cell in the E215Q mutant only ranged from zero to four. However, 84% of the E215Q mutant cells did not assemble any hooks. Importantly, these findings indicate that the reduced class 3 gene expression in the R177H and G183A mutants did not result from a reduced number of hooks or slower hook polymerization. The reduced number of hooks in the mutants E211Q and E215Q likely influenced the prolonged suppression of class 3 gene expression.

The simultaneous hook and filament labelling also served for determining the percentage of attached filaments per cell (**Fig. 3C**, **Fig. S4**). Therefore, filaments were counted on previously analyzed cells with determined hook numbers. Since a filament can only be assembled if the HBB formation is completed and the fT3SS has undergone the substrate specificity switch, all cells without HBBs were excluded from this analysis. We therefore reasoned that the frequency of filament formation, meaning the ratio of hooks with an attached filament, would be reduced in mutants with a defective substrate specificity switch. The distribution of attached filaments in the G183A mutant was similar to the WT distribution (**Fig. S4**), however overall percentage of hooks with attached filaments has decreased from 59% in the WT to 50% in G183A (**Fig. 3C**). Even though the R177H mutant did not have a defect in HBB assembly, the ratio of hooks with attached filaments was reduced to 38% in the mutant strain. In the ATPase activity mutants E211D and E211Q, the maximum number of filaments per cell was reduced to four and two, respectively. Consequently, the percentage of hooks with attached filaments decreased to 25% and 11%, respectively. Nonetheless, both mutants (E211D, E211Q) showed filament formation to some extent, whereas no filaments were detected in E215Q, independent from the number of assembled HBBs. In summary, the reduction of hooks with attached filaments in the FliI mutant strains support a defective substrate specificity switch.

To further confirm the involvement of the FliI ATPase in the substrate specificity switch, we measured the hook lengths of the WT and R177H and G183A mutants (**Fig. 3D-E**). We hypothesized that the delayed substrate specificity switch in the mutants would prolong the operating time of the T3SS in its early substrate secretion mode, resulting in elongated hooks. Therefore, we purified HBB complexes from the WT, and R177H and G183A mutants and measured the lengths of the hooks. Importantly, the lengths of the hooks of both mutants were slightly increased. The WT displayed hooks with an average length of 54.4 nm, whereas the hooks of the R177H mutant were on average 57.1 nm. In line with the increased secretion of early substrates (**Fig. 2C**) in the G183A mutant, the hooks from this mutant were even longer, on average 58.5 nm. The elongated hooks assembled by the FliI mutants further confirm an involvement of the fT3SS ATPase FliI in the substrate specificity switch.

Collectively, our results demonstrate that FliI couples ATPase activity to substrate specificity switching of the fT3SS. The introduced point mutations impaired the transition to late substrate secretion, as reflected in reduced class 3 gene expression, fewer hooks with attached filaments, and elongated hooks. These effects are not due to defects in FliI oligomerization or basal body localization, as confirmed by *in vivo* photo-crosslinking and fluorescence microscopy. Thus, while FliI ATPase activity is dispensable for early substrate secretion, it is important for efficient progression to the late assembly mode.

## Discussion

ATP hydrolysis by the FliI ATPase is essential for efficient operation of the fT3SS, but its precise contribution to the temporal regulation of flagellar assembly has remained unclear. In this study, we show that FliI activity is dispensable for early substrate secretion but critically required for the substrate specificity switch and efficient export of late substrates. The observed defect in the secretion of late substrates was not limited to FliC, as secretion of other late substrates (FlgM, FlgK, FliD) was similarly impaired (**Fig. S2C**), supporting a general role of FliI in facilitating late substrate export. Our analysis of FliI point mutants demonstrates that they exhibit impaired late-stage assembly, with active early substrate export and HBB formation, revealing a role for FliI in supporting the transition to late secretion.

The key findings are summarized in **Table 1**. Notably, mutants such as R177H and G183A, despite retaining early secretion and WT numbers of HBBs, exhibited strong defects in late substrate secretion and reduced gene expression from class 3 promoters. Mutants with impaired ATPase activity (E211D, E211Q, E215Q) showed progressively more severe defects in all late assembly metrics. Importantly, these defects were not due to loss of FliI oligomerization or localization, as demonstrated by *in vivo* photo-crosslinking and fluorescence microscopy. Together, our data suggest that the function of the ATPase activity of FliI is not merely to contribute to protein translocation but also facilitates the transition of the fT3SS to late substrate export.

**Table 1.** Characteristics of the analyzed FliI mutants.

| | Relative motility | Relative early substrate secretion | Relative FliC-HiBiT secretion | Relative class 3 gene expression $T_{90}$ | Hooks per cell | Hooks with attached filament [%] | Hook length [nm] |
| --- | --- | --- | --- | --- | --- | --- | --- |
| WT | 1.00 | 1.00 | 1.000 | 1.000 | 3.1 | 58.7 | 54.4 |
| $\Delta flil$ | 0.04 | 0.02 | 0.001 | 0.017 | NA | NA | NA |
| R177H | 0.48 | 1.00 | 0.417 | 0.183 | 3.1 | 38.2 | 57.1 |
| G183A | 0.93 | 1.38 | 0.754 | 0.778 | 3.0 | 50.5 | 58.5 |
| E211D | 0.25 | 0.62 | 0.135 | 0.034 | 2.6 | 24.9 | NA |
| E211Q | 0.09 | 0.37 | 0.003 | 0.027 | 0.5 | 11.4 | NA |
| E215Q | 0.06 | 0.13 | 0.001 | 0.022 | 0.1 | 0.0 | NA |

Recent structural and functional studies have proposed that FliHI modulate FlhA conformation to stabilize the late export state (47). Our findings are fully consistent with this model. It is likely that reduced ATPase activity or altered protein-protein interfaces in our mutants impair this remodeling function, thereby disrupting efficient switching to late substrate export. Additionally, we cannot exclude that FliI activity may contribute to dissociation of late chaperone-substrate complexes, as shown for the ATPase InvC of the evolutionary related virulence associated T3SS - processes that may be more ATP-dependent than early substrate export (26).

Taken together, our findings support a model in which FliI performs multiple, coordinated functions during flagellar assembly. First, FliI acts as a dynamic substrate carrier, delivering substrates to the export gate (24). Second, it serves as an energy-coupling ATPase that facilitates the efficient translocation of substrates through the fT3SS (27). Third, FliI plays an active role in controlling the substrate specificity switch, likely by modulating the conformational state of the export gate and promoting the transition from early to late substrate export (47). Hence, rather than serving solely as an energy-coupling translocase, FliI operates as a multi-functional ATPase that integrates substrate delivery, energy coupling, and specificity switching, underscoring its central role in flagellar biogenesis.\

## Materials and Methods

### Bacterial strains, plasmids and media

All bacterial strains and plasmids used in this study are listed in Table S1 and were derived from *S. enterica* subsp. *enterica* serovar Typhimurium LT2. Cells were cultivated at 37 °C under aerobic conditions in lysogeny broth (LB) at 180 rpm. If not stated otherwise, main cultures were diluted 1:100 and grown to mid-exponential phase. If required, cultures were supplemented with ampicillin (cf = 100 µg/mL), chloramphenicol (cf = 12.5 or 10 µg/mL) and/or AnTc (cf = 100 ng/mL). The FliI point mutations were introduced chromosomally by site-specific mutagenesis using λ-Red homologous recombination (48). Oligonucleotides used in this study are listed in Table S2. Furthermore, the generalized transducing phage of *Salmonella enterica* serovar Typhimurium P22 HT105/1 int-201 was used (49). Plasmids were constructed using NEBuilder® HiFi DNA Assembly Master Mix (New England Biolabs, catalogue number E2621L) ore one-step mutagenesis (50).

### Swimming motility assay

Swimming motility was studied using tryptone broth-based soft agar swim plates containing 0.3% Bacto agar. Motility plates were inoculated with 2 µL of overnight culture and incubated at 37 °C for 4-5 h. Images were acquired by scanning the plates and the diameters of the swimming halos were measured using Fiji (51). The swimming diameters of the mutant strains were normalized to those of the WT.

### Subcellular localization of FliI

The strains carried translational fusions of *fliG* to mNeonGreen and of HaloTag to *fliI*. Additionally, the strains carried an AnTc inducible P*_tetA_* promoter for the flagellar master regulator operon *flhDC* (32). Overnight cultures were grown in 2 mL LB. Cultures were diluted 1:100 in 10 mL of fresh LB and incubated at 37 °C for 1.5 h. The expression of *flhDC* was induced by the addition of AnTc (cf = 100 ng/µL), followed by 1 h incubation at 37 °C. A 500 µL sample of each culture was collected and centrifuged at 2,550 × g for 3 min. Pellets were resuspended in 1 × PBS and adjusted to an OD_600_ of 0.3. A 150 µL sample of each strain was transferred to a fresh 1.5 mL reaction tube and HaloTag labeling was performed by adding the HaloTag ligand tetramethylrhodamine (cf = 0.5 µM, Promega, catalogue number G8251) and incubating the samples for 30 min at 37 °C. Cells were washed twice with 500 µL of 1 × PBS and resuspended in 300 µL of 1 × PBS for microscopy. For imaging, 1 µL of labeled cells was spotted onto a 1% agarose pad in 1 × PBS. The cell suspension spot was air-dried for approximately 1 min and plasma-cleaned glass coverslips were applied. Fluorescence microscopy was performed using a Ti-2 Nikon inverted microscope equipped with a CFI Plan Apochromat DM 60× Lambda oil Ph3/1.40 (Nikon) oil objective, an Orca Fusion BT camera (Hamamatsu), and a SPECTRA III LED light source (Lumencor). Z-stacks were acquired in a 1.6 µm range with 5 stacks at 0.4 µm intervals. Microscopy data was analyzed using a custom Jupyter Notebook (See data and code availability statement) and Ilastik v1.4.0 (52).

### *In vivo* photo-crosslinking of the FliI hexamer

The protocol for *in vivo* photo-crosslinking to study the ability of FliI to form hexamers was adapted from previously published protocols (53, 54). Strains were deleted for *fliI*, carried pSUP and expressed FliI_3×FLAG_ (with amber mutation and respective point mutations) constitutively from pTrc99A-FF4. Overnight cultures were grown in LB, supplemented with chloramphenicol (cf = 10 µg/mL) and ampicillin (cf = 100 µg/mL) at 37 °C, 180 rpm. Main cultures were diluted 1:100 in 10 mL LB with the respective supplements in the presence of 1 mM *p*Bpa at 37 °C until they reached mid-exponential phase. 9 mL of each culture was harvested at 4,000 × g, 4 °C for 5 min. Cells were washed once with 2 mL cold 1 × PBS and pelleted again with centrifugation. Cells were resuspended in 2 mL cold 1 × PBS. 1 mL of the cell culture was transferred into a fresh 1.5 mL reaction tube; another 1 mL was transferred into a 6-well cell culture dish and irradiated for 30 min with UV light (λ = 366 nm) on ice. Afterwards, the cells were transferred into a fresh 1.5 mL reaction tube harvested at 10,000 × g at 4 °C for 3 min. Proteins were precipitated using 10% (v/v) TCA, followed by 30 min incubation on ice. Proteins were pelleted by centrifugation at 20,000 × g for 30 min at 4 °C. Protein pellets were washed with ice-cold acetone and air-dried. Samples were adjusted to 50 OD units/µL. 500 OD units were analyzed under denaturing conditions using SDS-PAGE. Immunoblotting was performed using primary α-FLAG antibody (Sigma-Aldrich, catalogue number F1804-200UG, 1:10,000 in 1 × TBS-T). Proteins were detected using secondary α-mouse antibodies conjugated to horseradish peroxidase (Bio-Rad Immun-Star Goat Anti-Mouse (GAM)-HRP Conjugate, catalog number 170-5047, 1:20,000 in 1 × TBS-T).

### Minimal inhibitory concentration assay (FlgE-Bla secretion)

The minimal inhibitory concentration (MIC) of ampicillin correlating with the amount of secreted hook-protein-β-lactamase fusion protein (FlgE-Bla) was determined as described by Lee et al. (36) and adapted by Singer et al. (37) to fit a 96-well plate format. Briefly, cultures were grown overnight in 200 µL LB in a 96-well plate at 37 °C, 180 rpm in biological triplicates. The cultures were diluted 1:100 in fresh LB and grown to mid-exponential phase at 37 °C at 180 rpm. The cultures were then diluted 1:10 in 300 µL LB and finally diluted 1:10 in 200 µL LB media supplemented with increasing concentrations of fresh ampicillin (Amp) in technical triplicates. Cells were grown for 4.5 h at 37 °C at 180 rpm and OD_600_ was measured with a microplate reader. The IC_50_ was calculated using a custom Jupyter Notebook.

### Secretion Assay using the split NanoBit® enzyme

Nano-Glo® HiBiT Extracellular Detection System was purchased from Promega (N2420). Quantification of the secreted HiBiT-tagged proteins was determined as described previously (41). Briefly, overnight cultures were diluted 1:100 and grown to late exponential phase at 37 °C at 180 rpm. OD_600_ was measured and 1 mL sample of each culture was transferred into a 1.5 mL Eppendorf tube on ice. Samples were centrifuged at 13,000 x g at 4 °C for 3 min. 500 µL of the supernatant (SN) was transferred into a fresh Eppendorf tube and stored on ice. 25 µL of the SN was transferred in technical triplicates into a white 384-well plate with LB-blanks included. The working solution was prepared and SN samples and working solution were mixed as described by the manufacturer. Luminescence was measured at T_0_ = 0 min and T_10_ = 10 min using a microplate reader. The relative light units (RLU) of each strain were calculated using following equation:

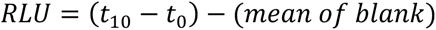

The calculated values were normalized to the measured OD_600_ and to the WT strain.

### Secretion Assay analyzed by SDS-PAGE and Western Blot

For analysis with SDS-PAGE and Western Blotting, the same cultures were used as for the secretion assay using the split luciferase. A 1.9 mL aliquot was taken, and cells were harvested by centrifugation at 13,000 × g for 5 min. 1 mL of the supernatant was transferred into a fresh tube, the remaining supernatant was discarded, and the cell pellets were resuspended in 1 mL of double-distilled water. Proteins were precipitated using 10% (v/v) TCA, followed by 30 min incubation on ice. Proteins were pelleted by centrifugation at 20,000 × g for 30 min at 4 °C. Protein pellets were washed with ice-cold acetone and air-dried. Samples were adjusted to 20 OD units/µL. 200 OD units were analyzed under denaturing conditions using SDS-PAGE. The samples of all three biological replicates were pooled to obtain the mean protein amount of all replicates. Immunoblotting was performed using primary α-FlgM (gift from K. Hughes, 1:10,000 in 1 × TBS-T), α-FlgK (gift from T. Minamino, 1:10,000 in 1 × TBS-T), α-FliD (gift from T. Minamino, 1:10,000 in 1 × TBS-T) or α-FliC (Difco, catalog number 228241 Salmonella H Antiserum I, 1:5,000 in 1 × TBS-T) antibodies. Proteins were detected using secondary α-rabbit antibodies conjugated to horseradish peroxidase (Bio-Rad Immun-Star Goat Anti-Rabbit (GAR)-HRP Conjugate, catalog number 170-5046, 1:20,000 in 1 × TBS-T). The relative amounts of secreted and cellular proteins were determined by normalization to the housekeeping protein DnaK using the Image Lab software (Bio-Rad). DnaK was detected using a primary α-DnaK (abcam, catalog number ab69617, 1:10,000 in 1 × TBS-T) antibody and secondary α-mouse antibodies conjugated to horseradish peroxidase (Bio-Rad Immun-Star Goat Anti-Mouse (GAM)-HRP Conjugate, catalog number 170-5047, 1:20,000 in 1 × TBS-T).

### Luciferase assay – P*_motA_*-*luxCDABE*

Strains harboring the plasmid pRG19 (P*_motA_*-*luxCDABE*) were used to monitor flagellar class 3 gene expression (44). ON-cultures of 6 biological replicates were grown in 200 µL LB supplemented with Chloramphenicol (Cm) with cf = 12.5 µg/ml in a 96-well plate at 37 °C, 200 rpm. To prevent evaporation, the plate was covered with a Breath-Easy sealing membrane. The cultures were diluted 1:10 in 200 µL LB supplemented with Cm and again 1:10 in LB supplemented with Cm and AnTc to a final concentration of 100 ng/ml to induce expression of the flagellar master regulator from P*_tetA_* (32) in a white 96-well plate for final dilution to 1:100. The OD_600_ was measured, as well as luminescence with a microplate reader for 6 h every 15 min. The relative light units (RLU) were normalized to the measured OD_600_ of each sample and analyzed as described previously (45, 55). The relative class 3 gene expression in respect to the WT was calculated for three timepoints T_360_ (360 min).

### Hook and Filament Staining

For determination of the number of hooks per cell and the number of hooks with an attached filament, cells were grown to mid-log phase. 500 µL samples were taken and centrifuged at 2500 × g for 5 min. The cells were carefully resuspended in 500 µL fresh 1 × PBS. Star Green maleimide dye (Abberior, catalogue number STGREEN-0003-1MG) with cf = 10 µM and 0.5 µL α-HA-Alexa555 (Thermo Fisher Scientific, catalogue number 26183-A555) were added to stain the filaments (*fliC*_T237C_) (56) with maleimide and the hook (*flgE*_3×HA_) with immunostaining. The cells were incubated for 1 h at 37 °C at 300 rpm. The cells were washed twice with 1 × PBS before loading the samples on a poly-L-lysine coated coverslip as described previously (57, 58). The cells were fixed using 2% paraformaldehyde-0.2% glutaraldehyde solution and rinsed with 1 × PBS before staining the DNA fluoroshield containing DAPI (Sigma Aldrich, catalogue number F6057-20ML). Cells were observed using a Zeiss Inverted Microscope Axio Observer Z1 at 100 × magnification and z-stack experiments were performed with 9 slices, 4 µm range with 0.5 µm slice interval. Analysis was done using Fiji for image processing equipped with the MicrobeJ Plugin to determine the number of hooks per cell (51, 59). Filaments were counted manually on cells with determined hook numbers.

### Hook Length Measurements

The purification of hook basal body complexes was adapted from a previously published protocol (6, 12, 60). Briefly, an overnight culture was inoculated with a single colony in LB medium. The next day, 500 mL LB medium was inoculated 1:100 with the overnight culture and grown for 2.5 h. We utilized strains with an AnTc inducible P*_tetA_* promoter for the flagellar master regulator operon *flhDC* (32). To induce flagella biosynthesis, AnTc was added, and the cells were further inoculated for 1 h. Cells were harvested at 4,000 × g at 4 °C for 15 min. The cell pellet was carefully resuspended in 20 mL ice-cold sucrose solution (0.5 M sucrose, 0.15 M Trizma base, unaltered pH) on ice. Lysozyme and EDTA pH 4.7 were added slowly to final concentrations of 0.1 mg/mL and 2 mM, respectively, while stirring the cell suspension on ice. After 5 min of stirring on ice, the suspension was transferred to RT and slowly stirred for 1 h to allow spheroplast formation. For cell lysis, Triton X-100 was added to a final concentration of 1%. After rapidly stirring the suspension for 10 min until it became translucent, it was slowly stirred further for 30-45 min on ice. To degrade DNA, 2 mg DNAseI and MgSO_4_ were added to a final concentration of 5 mM while stirring on ice. After 5 min, EDTA pH 4.7 was added at a final concentration of 5 mM. To pellet cell debris and unlysed cells, the suspension was centrifuged at 15,000 × g at 4 °C for 10 min. The supernatant was collected and centrifuged at 37,000 rpm (T-647.5 Fixed Angle Rotor, Thermo Fisher Scientific) for 1 h at 4 °C to pellet the flagella. The flagella were washed carefully with 50 mL buffer A (0.1 M KCl, 0.3 M sucrose, 0.1% Triton X-100) and collected again at 37,000 rpm (T-647.5 Fixed Angle Rotor, Thermo Fisher Scientific) for 1 h at 4 °C. To depolymerize the flagellin filament, the pellet was carefully resuspended in 50 mL buffer B (50 mM glycine, 0.1% Triton X-100, pH 2.5), followed by 30 min incubation at room temperature. Finally, the hook basal bodies were collected again by centrifugation at 37,000 rpm (T-647.5 Fixed Angle Rotor, Thermo Fisher Scientific) for 1 h at 4 °C. Hook basal bodies were resuspended carefully in 50 µL buffer C (10 mM Tris pH 8, 5 mM EDTA pH 8, 0.1% Triton X-100) and incubated overnight on a rolling platform at 4 °C. Grids were prepared as described previously (61). Briefly, aliquots of HBB samples were applied to freshly glow discharged carbon-film-coated copper grids and allowed to adsorb for 10 minutes. After three washes with distilled water, the grids were contrasted with 4% phospho-tungstic-acid/1% trehalose, touched on filter paper and air-dried. The grids were analyzed in a Leo 906 transmission electron microscope (Zeiss) at 100 kV acceleration voltage. Micrographs were scanned using a Morada side-mount camera (SIS) with ImageSP software from TRS (Tröndle). Hook lengths were measured using Fiji (51).

### Statistical Analyses

Statistical analyses were performed using GraphPad Prism 10 (GraphPad Software), and values of P < 0.05 were considered statistically significant.

## Data and code availability

The Jupyter notebook used for colocalization analyses of the microscopy data is available at https://github.com/SalmoLab/FliI_ATPase.

## Acknowledgements

M.E. acknowledges funding from the Deutsche Forschungsgemeinschaft (DFG) research grant no. 322866343 and from the Max Planck Society as Max Planck Fellow. M.D.G. was supported by a Grand Challenge Initiative Global Health grant of the Berlin University Alliance (no.113_MC_GH_MEL-BER_Erhardt_HU). We thank K. Hughes (The University of Utah) and T. Minamino (Osaka University) for providing antibodies. We thank members of the Erhardt lab for helpful discussions and critical comments on the manuscript, and Heidi Landmesser and Raúl Trepel-Washington for technical assistance.

## Author Contributions

M.E. conceptualized and supervised the research project and ensured funding. R.E. and M.D.G. generated *S. enterica* mutants. R.E. performed the experiments, wrote the first draft of the manuscript and prepared figures. L.H. wrote the code for colocalization analyses for the microscopy data. C.G. prepared TEM grids and observed the purified HBB samples. M.E. reviewed and edited the manuscript. All authors reviewed the results and approved the final version of the manuscript.

