## Supplementary Information for "The FliI ATPase couples ATP hydrolysis to substrate switching in bacterial flagellar type-III secretion"

Running Head: FliI ATPase Controls Flagellar Secretion Specificity

**Table of Contents**

Figure S1. Localization and oligomerization of FliI and FliI mutants.
Figure S2. FliI is required for efficient secretion of late flagellar substrates.
Figure S3. Quantification of class 3 gene expression.
Figure S4. FliI mutants exhibit a decreased number of hooks with attached filaments.

Table S1. Strains and plasmids used in this study.
Table S2. Oligonucleotides used in this study.

Supplementary References

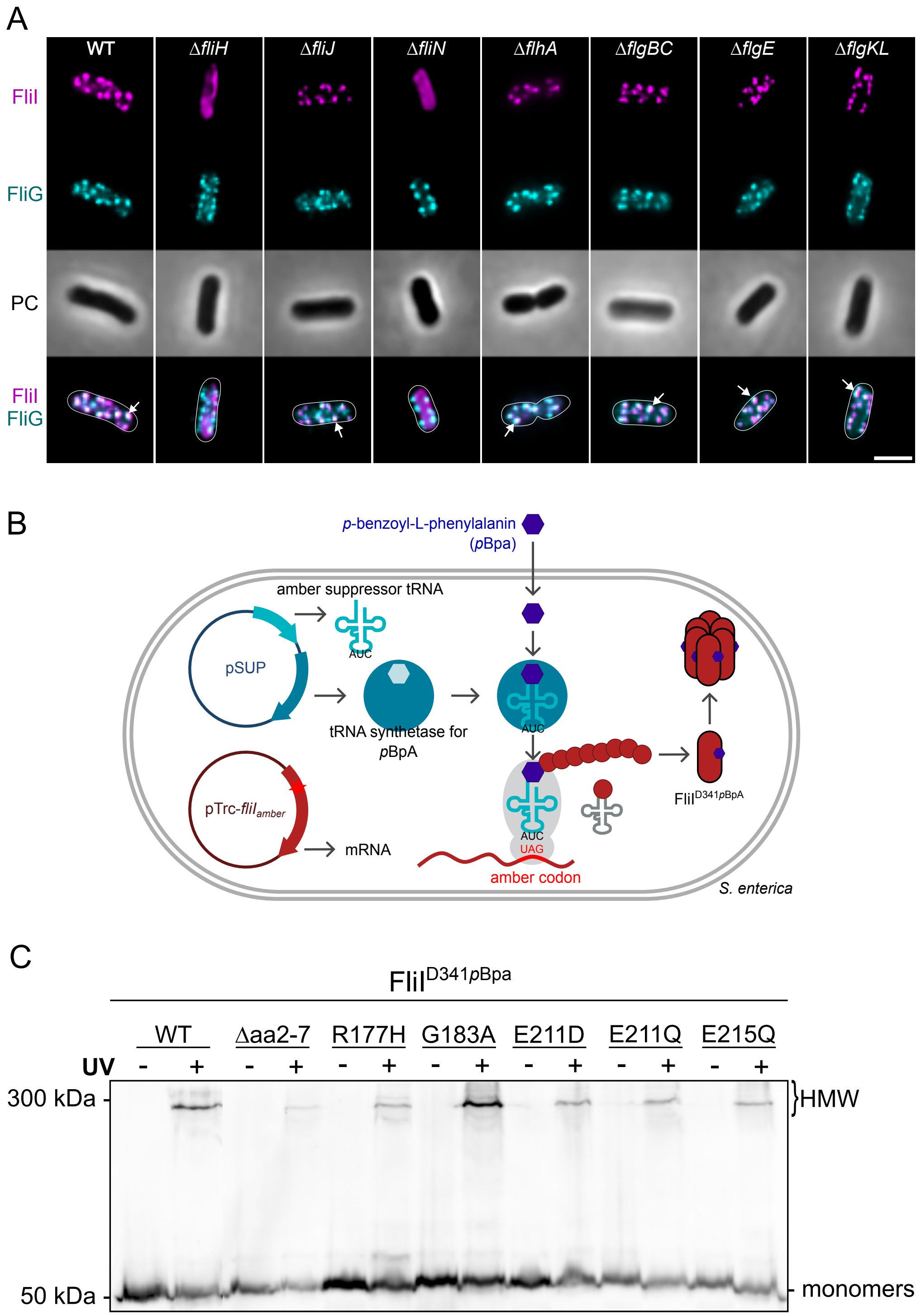


**Fig. S1**. Localization and oligomerization of FliI and FliI mutants. (A) Representative microscopy images of FliI-HaloTag in various genetic backgrounds. FliI-HaloTag labeled with TMR-ligand (magenta), basal bodies visualized with mNeonGreen-FliG (cyan); PC (PhaseContrast). Scale-bar: 2 µm. (B) Schematic of the *in vivo* site-specific photo-crosslinking system used to assess FliI oligomerization. An amber codon is introduced at position D341 of FliI to incorporate the photo-reactive amino acid p-benzoyl-L-phenylalanine (pBpa). Upon UV irradiation, crosslinking occurs between interacting FliI subunits. (C) *In vivo* UV crosslinking of FLAG-tagged FliI$^{D341pBpa}$ variants. High molecular weight (HMW) adducts were detected for WT and all point mutants, confirming oligomerization. Samples were separated by SDS-PAGE and visualized by immunoblotting against FLAG.

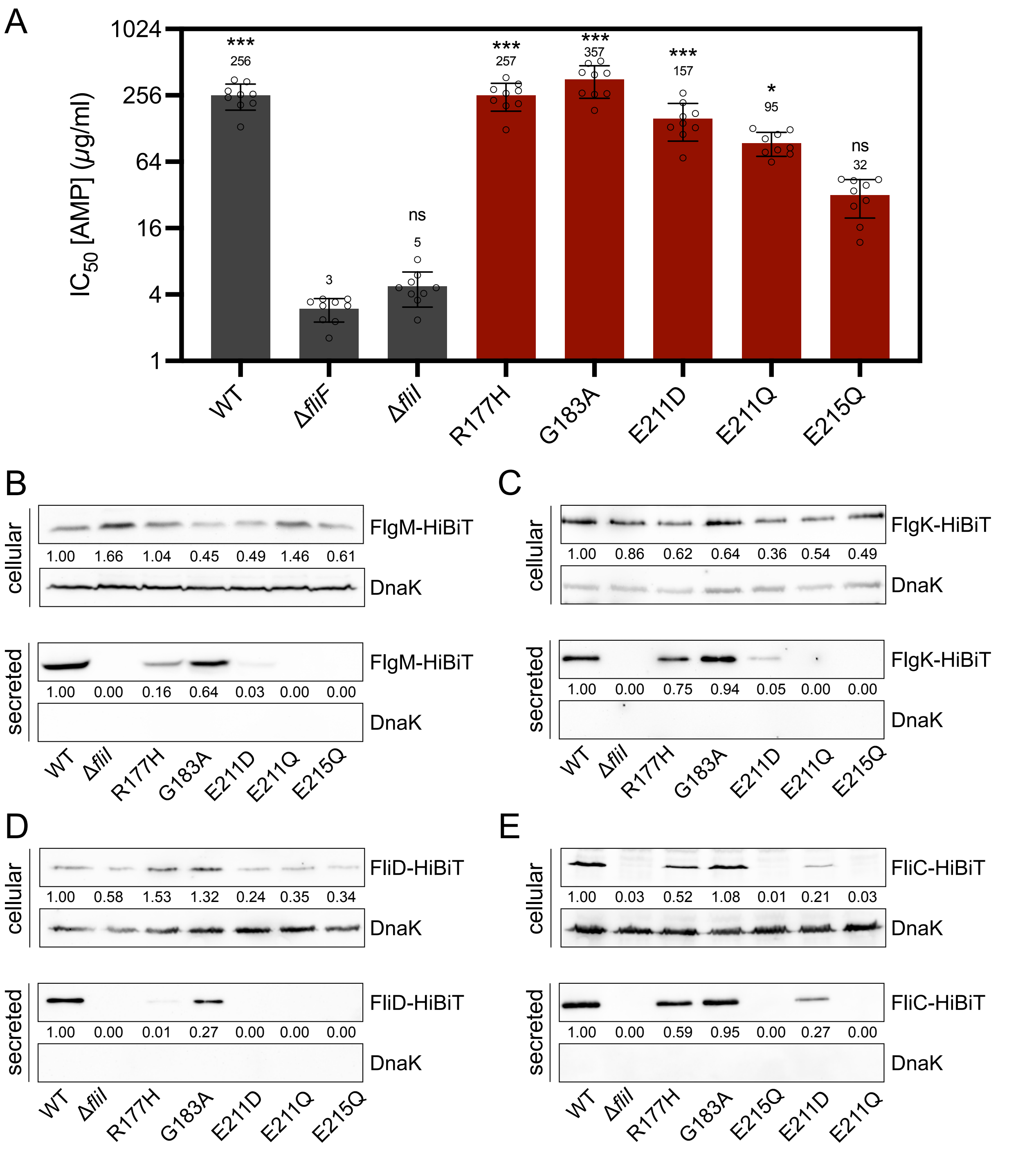


**Fig. S2**. FliI is required for efficient secretion of late flagellar substrates. (A) Quantification of early substrate (FlgE-Bla) secretion. Shown are $IC_{50}$ values for ampicillin resistance, reflecting the amount of secreted FlgE-Bla. Bars represent means ± SD with individual data points shown from ≥3 independent experiments. Statistical significance was determined by one-way ANOVA with Dunnett's multiple comparisons test vs. Δ*fliF.* ***p < 0.001; **p < 0.01; *p < 0.05; ns, not significant. (B-E) Western Blot analysis of FlgM-HiBiT (B), FlgK-HiBiT (C), FliD-HiBiT (D) and FliC-HiBiT in cellular and secrete fractions. Samples were pooled from 3 independent experiments, separated by SDS-PAGE and immunoblotted using anti-DnaK and anti-FlgM, anti-FlgK, anti-FliD or anti-FliC, respectively. Numbers indicate the amount of HiBiT-tagged proteins normalized to DnaK and to the WT.

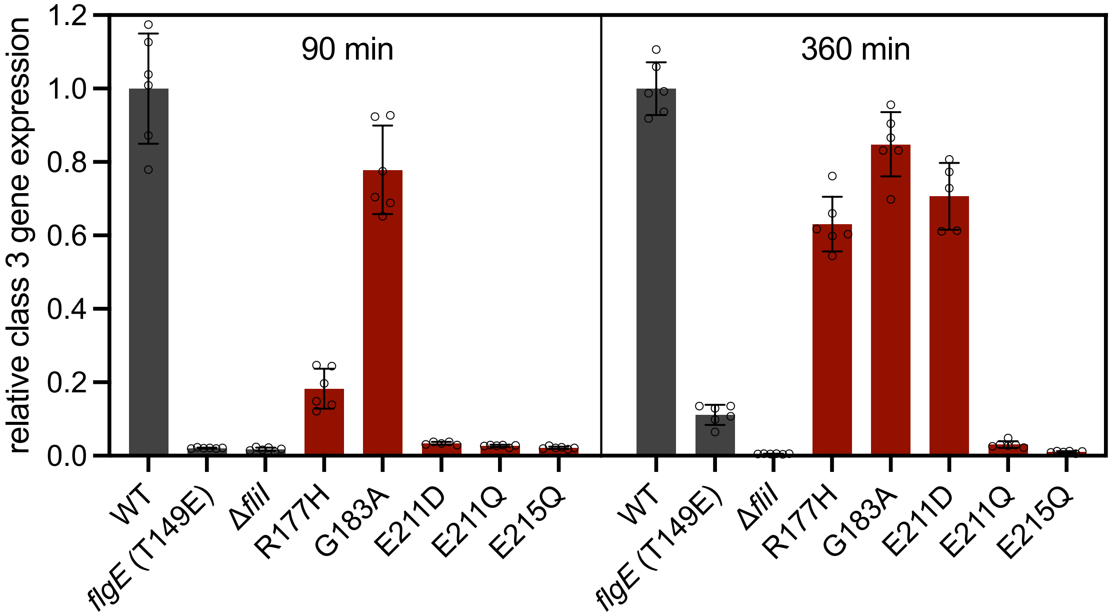


**Fig. S3**. Quantification of class 3 gene expression. Relative class 3 gene expression ($P_{motA}$ activity) at 90 min ($T_{90}$) and 360 min ($T_{360}$) after induction of flagellar synthesis. Shown are the normalized luminescence values from six biological replicates, expressed relative to the WT signal at each time point. Bars represent means ± SD with individual data points shown.

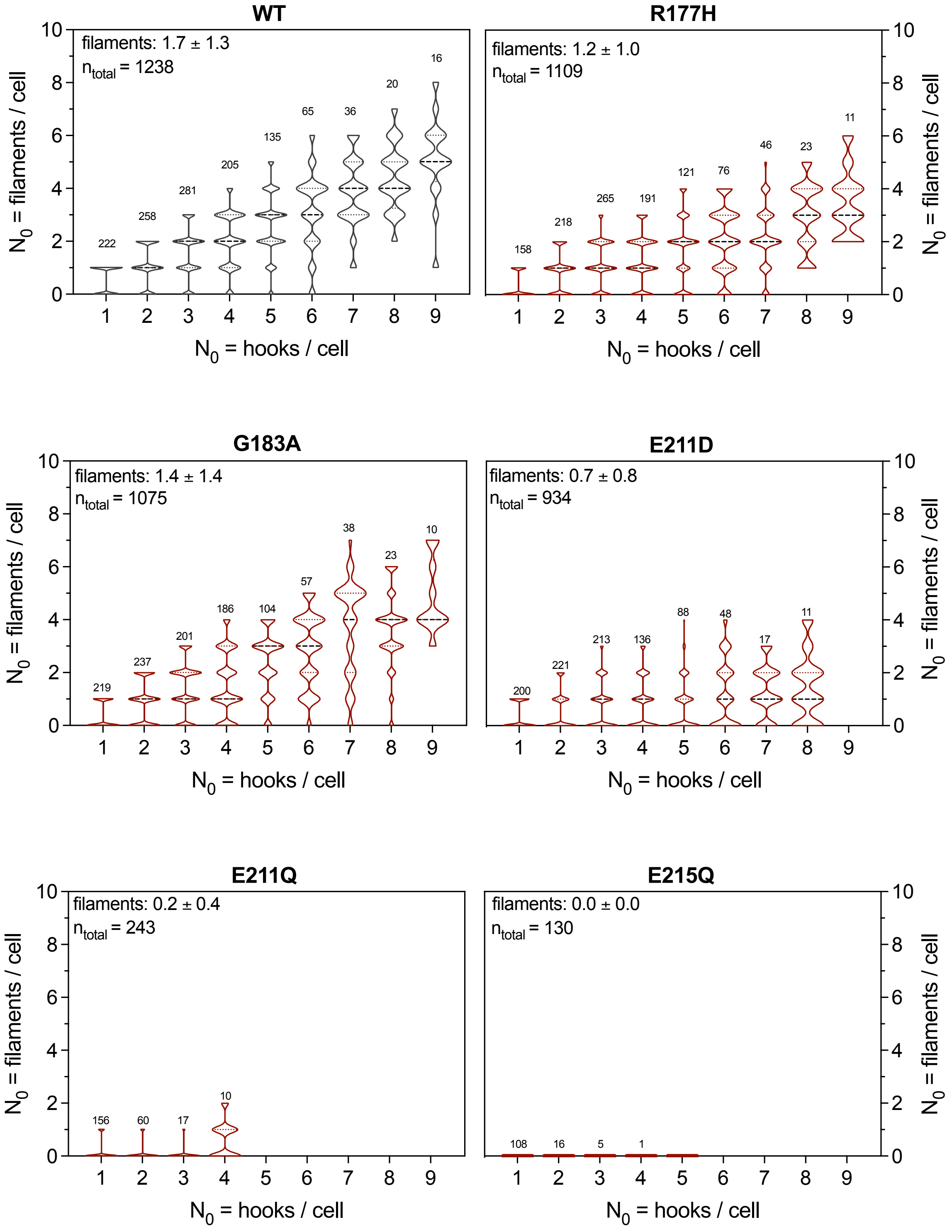


**Fig. S4**. FliI mutants exhibit a decreased number of hooks with attached filaments. Shown are the numbers of filaments per cell as a function of hook number per cell. Cells were FliC$^{ON}$ and expressed *flgE*$_{3\times HA}$ and *fliC*$_{T237C}$ for immuno- and maleimide staining of the hook and filament, respectively. The width of the violin plot outline illustrates the distribution of the data. The numbers above each violin bar represent the number of counted cells for this category. Thick dashed lines represent the median; dotted lines represent the quartiles.

**Table S1.** Strains and Plasmids used in this study.

| Strain | Genotype | Source/Reference |
|---|---|---|
| EM2624 | LT2 wild-type | Lab Collection |
| TH12424 | Δ*fliI*7364 | Lab Collection |
| EM4120 | *fliI*23023 (R177H) | Lab Collection |
| EM4121 | *fliI*23024 (G183A) | Lab Collection |
| EM9307 | *fliI*23209 (E211D) | Lab Collection |
| EM9308 | *fliI*23210 (E211Q) | Lab Collection |
| EM4122 | *fliI*23025 (E215Q) | Lab Collection |
| EM14153 | *fliI*23109 (*fliI*-HaloTag) *fliG*22799 (mNeonGreen-*fliG*) $P_{flhDC}$5451::Tn10dTc[del-25] | This study |
| EM14391 | *fliG*22799 (mNeonGreen-*fliG*) $P_{flhDC}$5451::Tn10dTc[del-25] *fliI*23619 (R177H)::HaloTag (C-term) | This study |
| EM15105 | *fliG*22799 (mNeonGreen-*fliG*) $P_{flhDC}$5451::Tn10dTc[del-25] *fliI*236677 (G183A)::HaloTag (C-term) | This study |
| EM14392 | *fliG*22799 (mNeonGreen-*fliG*) $P_{flhDC}$5451::Tn10dTc[del-25] *fliI*23620 (E211D)::HaloTag (C-term) | This study |
| EM14393 | *fliG*22799 (mNeonGreen-*fliG*) $P_{flhDC}$5451::Tn10dTc[del-25] *fliI*23621 (E211Q)::HaloTag (C-term) | This study |
| EM14394 | *fliG*22799 (mNeonGreen-*fliG*) $P_{flhDC}$5451::Tn10dTc[del-25] *fliI*23622 (E215Q)::HaloTag (C-term) | This study |
| EM15149 | *fliG*22799 (mNeonGreen-*fliG*) $P_{flhDC}$5451::Tn10dTc[del-25] *fliI*23681(Δaa2-7)::HaloTag (C-term) | This study |
| EM14389 | *fliI*23109 (*fliI*-HaloTag) *fliG*22799 (mNeonGreen-*fliG*) $P_{flhDC}$5451::Tn10dTc[del-25] Δ*fliH*7363 | This study |
| EM14390 | *fliI*23109 (*fliI*-HaloTag) *fliG*22799 (mNeonGreen-*fliG*) $P_{flhDC}$5451::Tn10dTc[del-25] Δ*fliJ*7365 | This study |
| EM15298 | *fliI*23109 (*fliI*-HaloTag) *fliG*22799 (mNeonGreen-*fliG*) $P_{flhDC}$5451::Tn10dTc[del-25] Δ*fliN*23676::FRT | This study |
| EM15299 | *fliI*23109 (*fliI*-HaloTag) *fliG*22799 (mNeonGreen-*fliG*) $P_{flhDC}$5451::Tn10dTc[del-25] Δ*flhA*23464::FRT | This study |
| EM14428 | *fliI*23109 (*fliI*-HaloTag) *fliG*22799 (mNeonGreen-*fliG*) $P_{flhDC}$5451::Tn10dTc[del-25] Δ*flgBC*23587::FRT | This study |
| EM14429 | *fliI*23109 (*fliI*-HaloTag) *fliG*22799 (mNeonGreen-*fliG*) $P_{flhDC}$5451::Tn10dTc[del-25] Δ*flgE*22964::FRT | This study |
| EM14430 | *fliI*23109 (*fliI*-HaloTag) *fliG*22799 (mNeonGreen-*fliG*) $P_{flhDC}$5451::Tn10dTc[del-25] Δ*flgKL*5739::FRT | This study |
| EM14456 | Δ*fliI*7364 / pEM14338 (pTrc99A-FF4-*fliI* (D341amber)-SAGASA-3×FLAG, C-ter, Amp$^R$) / pSUP (Cm$^R$) | This study |
| EM15252 | Δ*fliI*7364 / pEM15178 (pTrc99A-FF4-*fliI* (Δaa2-7, D341amber)-SAGASA-3×FLAG, C-ter, Amp$^R$) / pSUP (artificial amino-acid, Cm$^R$) | This study |
| EM15253 | Δ*fliI*7364 / pEM15179 (pTrc99A-FF4-*fliI* (R177H, D341amber)-SAGASA-3×FLAG, C-ter, Amp$^R$) / pSUP (artificial amino-acid, Cm$^R$) | This study |
| EM15254 | Δ*fliI*7364 / pEM15180 (pTrc99A-FF4-*fliI* (G183A, D341amber)-SAGASA-3×FLAG, C-ter, Amp$^R$) / pSUP (artificial amino-acid, Cm$^R$) | This study |
| EM15255 | Δ*fliI*7364 / pEM15181 (pTrc99A-FF4-*fliI* (E211D, D341amber)-SAGASA-3×FLAG, C-ter, Amp$^R$) / pSUP (artificial amino-acid, Cm$^R$) | This study |

| | | |
|---|---|---|
| EM15256 | Δ*fliI*7364 / pEM15182 (pTrc99A-FF4-*fliI* (E211Q, D341amber)-SAGASA-3×FLAG, C-ter, $Amp^R$) / pSUP (artificial amino-acid, $Cm^R$) | This study |
| EM15257 | Δ*fliI*7364 / pEM15183 (pTrc99A-FF4-*fliI* (E215Q, D341amber)-SAGASA-3×FLAG, C-ter, $Amp^R$) / pSUP (artificial amino-acid, $Cm^R$) | This study |
| TH9949 | *flgE*6569::*bla* Δ*flgBC*6557 ($Amp^R$) | (1) |
| TH12465 | *flgE*6569::*bla* Δ*flgBC*6557 Δ*fliF*7387 | (2) |
| TH12473 | *flgE*6569::*bla* Δ*flgBC*6557 Δ*fliI*7395 | (2) |
| EM7801 | *flgE*6569::*bla* Δ*flgBC*6557 *fliI*23023 (R177H) | Lab Collection |
| EM7802 | *flgE*6569::*bla* Δ*flgBC*6557 *fliI*23024 (G183A) | Lab Collection |
| EM7803 | *flgE*6569::*bla* Δ*flgBC*6557 *fliI*23025 (E215Q) | Lab Collection |
| EM9432 | *flgE*6569::*bla* Δ*flgBC*6557 *fliI*23209 (E211D) | Lab Collection |
| EM9433 | *flgE*6569::*bla* Δ*flgBC*6557 *fliI*23210 (E211Q) | Lab Collection |
| EM9743 | *flgM*23271::HiBiT (C-ter, RBS FlgN duplicated) | Lab Collection |
| EM10103 | *flgM*23271::HiBiT (C-ter, RBS FlgN duplicated) Δ*fliI*7364 | This study |
| EM10104 | *flgM*23271::HiBiT (C-ter, RBS FlgN duplicated) *fliI*23023 (R177H) | This study |
| EM10105 | *flgM*23271::HiBiT (C-ter, RBS FlgN duplicated) *fliI*23024 (G183A) | This study |
| EM10107 | *flgM*23271::HiBiT (C-ter, RBS FlgN duplicated) *fliI*23209 (E211D) | This study |
| EM10108 | *flgM*23271::HiBiT (C-ter, RBS FlgN duplicated) *fliI*23210 (E211Q) | This study |
| EM18068 | *flgM*23271::HiBiT (C-ter, RBS FlgN duplicated) *fliI*23025 (E215Q) | This study |
| EM9999 | Δ*hin*-5717::FRT *flgK*23312::HiBiT (after aa442, 3×SAGASA-HiBiT-3×SAGASA) | Lab Collection |
| EM18069 | Δ*hin*-5717::FRT *flgK*23312::HiBiT (after aa442, 3×SAGASA-HiBiT-3×SAGASA) Δ*fliI*7364 | This study |
| EM18070 | Δ*hin*-5717::FRT *flgK*23312::HiBiT (after aa442, 3×SAGASA-HiBiT-3×SAGASA) *fliI*23023 (R177H) | This study |
| EM18071 | Δ*hin*-5717::FRT *flgK*23312::HiBiT (after aa442, 3×SAGASA-HiBiT-3×SAGASA) *fliI*23024 (G183A) | This study |
| EM18072 | Δ*hin*-5717::FRT *flgK*23312::HiBiT (after aa442, 3×SAGASA-HiBiT-3×SAGASA) *fliI*23209 (E211D) | This study |
| EM18073 | Δ*hin*-5717::FRT *flgK*23312::HiBiT (after aa442, 3×SAGASA-HiBiT-3×SAGASA) *fliI*23210 (E211Q) | This study |
| EM18074 | Δ*hin*-5717::FRT *flgK*23312::HiBiT (after aa442, 3×SAGASA-HiBiT-3×SAGASA) *fliI*23025 (E215Q) | This study |
| EM18085 | Δ*hin*-5717::FRT *fliD*23314::HiBiT (Δaa205-209, 3×SAGASA-HiBiT-3×SAGASA) Δ*flgKL*5739::FRT | This study |
| EM18174 | Δ*hin*-5717::FRT *fliD*23314::HiBiT (Δaa205-209, 3×SAGASA-HiBiT-3×SAGASA) Δ*flgKL*5739::FRT Δ*fliI*7364 | This study |
| EM18175 | Δ*hin*-5717::FRT *fliD*23314::HiBiT (Δaa205-209, 3×SAGASA-HiBiT-3×SAGASA) Δ*flgKL*5739::FRT *fliI*23023 (R177H) | This study |
| EM18176 | Δ*hin*-5717::FRT *fliD*23314::HiBiT (Δaa205-209, 3×SAGASA-HiBiT-3×SAGASA) Δ*flgKL*5739::FRT *fliI*23024 (G183A) | This study |
| EM18177 | Δ*hin*-5717::FRT *fliD*23314::HiBiT (Δaa205-209, 3×SAGASA-HiBiT-3×SAGASA) Δ*flgKL*5739::FRT *fliI*23209 (E211D) | This study |
| EM18178 | Δ*hin*-5717::FRT *fliD*23314::HiBiT (Δaa205-209, 3×SAGASA-HiBiT-3×SAGASA) Δ*flgKL*5739::FRT *fliI*23210 (E211Q) | This study |

| | | |
|---|---|---|
| EM18179 | Δ*hin*-5717::FRT *fliD*23314::HiBiT (Δaa205-209, 3×SAGASA-HiBiT-3×SAGASA) Δ*flgKL*5739::FRT *fliI*23025 (E215Q) | This study |
| EM10744 | Δ*hin*-5717::FRT *fliC*23299::HiBiT (Δaa201-213 3×SAGASA-HiBiT-3×SAGASA) Δ*flgKL*5739::FKF | (3) |
| EM10852 | Δ*hin*-5717::FRT *fliC*23299::HiBiT (Δaa201-213 3×SAGASA-HiBiT-3×SAGASA) *fliI*23023 (R177H) Δ*flgKL*5739::FKF | This study |
| EM10853 | Δ*hin*-5717::FRT *fliC*23299::HiBiT (Δaa201-213 3×SAGASA-HiBiT-3×SAGASA) *fliI*23024 (G183A) Δ*flgKL*5739::FKF | This study |
| EM10854 | Δ*hin*-5717::FRT *fliC*23299::HiBiT (Δaa201-213 3×SAGASA-HiBiT-3×SAGASA) *fliI*23209 (E211D) Δ*flgKL*5739::FKF | This study |
| EM10950 | Δ*hin*-5717::FRT *fliC*23299::HiBiT (Δaa201-213 3×SAGASA-HiBiT-3×SAGASA) Δ*fliI*7364 Δ*flgKL*5739::FKF | This study |
| EM10951 | Δ*hin*-5717::FRT *fliC*23299::HiBiT (Δaa201-213 3×SAGASA-HiBiT-3×SAGASA) *fliI*23025 (E215Q) Δ*flgKL*5739::FKF | This study |
| EM10952 | Δ*hin*-5717::FRT *fliC*23299::HiBiT (Δaa201-213 3×SAGASA-HiBiT-3×SAGASA) *fliI*23210 (E211Q) Δ*flgKL*5739::FKF | This study |
| TH5633 | $P_{flhDC}$5451::Tn10dTc[del-25] / pRG19 ($P_{motA}$-*luxCDABE*, $Cm^R$, $Tet^R$) | (4) |
| TH14117 | $P_{flhDC}$5451::Tn10dTc[del-25] *flgE*2219(T149N) / pRG19 ($P_{motA}$-*luxCDABE*, $Cm^R$, $Tet^R$) | (4) |
| EM9611 | Δ*fliI*7364 $P_{flhDC}$5451::Tn10dTc[del-25] / pRG19 ($P_{motA}$-*luxCDABE*, $Cm^R$, $Tet^R$) | This study |
| EM9612 | *fliI*23023 (R177H) $P_{flhDC}$5451::Tn10dTc[del-25] / pRG19 ($P_{motA}$-*luxCDABE*, $Cm^R$, $Tet^R$) | This study |
| EM9613 | *fliI*23024 (G183A) $P_{flhDC}$5451::Tn10dTc[del-25] / pRG19 ($P_{motA}$-*luxCDABE*, $Cm^R$, $Tet^R$) | This study |
| EM9614 | *fliI*23025 (E215Q) $P_{flhDC}$5451::Tn10dTc[del-25] / pRG19 ($P_{motA}$-*luxCDABE*, $Cm^R$, $Tet^R$) | This study |
| EM11049 | *fliI*23209 (E211D) $P_{flhDC}$5451::Tn10dTc[del-25] / pRG19 ($P_{motA}$-*luxCDABE*, $Cm^R$, $Tet^R$) | This study |
| EM11050 | *fliI*23210 (E211Q) $P_{flhDC}$5451::Tn10dTc[del-25] / pRG19 ($P_{motA}$-*luxCDABE*, $Cm^R$, $Tet^R$) | This study |
| EM9031 | Δ*hin*-5717::FRT *fliC*6500(T237C) *flgE*7742::3×HA (after aa241) | Lab collection |
| EM12038 | *fliI*23023 (R177H) Δ*hin*-5717::FRT *fliC*6500(T237C) *flgE*7742::3×HA (after aa241) | This study |
| EM9688 | *fliI*23024 (G183A) Δ*hin*-5717::FRT *fliC*6500(T237C) *flgE*7742::3×HA (after aa241) | This study |
| EM11123 | Δ*hin*-5717::FRT *fliC*6500(T237C) *flgE*7742::3×HA (after aa241) *fliI*23025 (E215Q) | This study |
| EM11124 | Δ*hin*-5717::FRT *fliC*6500(T237C) *flgE*7742::3×HA (after aa241) *fliI*23209 (E211D) | This study |
| EM11125 | Δ*hin*-5717::FRT *fliC*6500(T237C) *flgE*7742::3×HA (after aa241) *fliI*23210 (E211Q) | This study |
| TH3730 | $P_{flhDC}$5451::Tn10dTc[del-25] | (5) |
| EM9565 | *fliI*23023 (R177H) $P_{flhDC}$5451::Tn10dTc[del-25] | This study |
| EM9566 | *fliI*23024 (G183A) $P_{flhDC}$5451::Tn10dTc[del-25] | This study |
| **Plasmid** | **Genotype** | **Source** |

| | | |
|---|---|---|
| pRG19 | $P_{motA}$-*luxCDABE*, $Cm^R$, $Tet^R$ | (6) |
| pSUP | $P_{glnS}$-BpaRS, $P_{proK}$-6TRN, $Cm^R$ | (7) |
| pEM14338 | pTrc99A-FF4-*fliI* (D341amber)-SAGASA-3×FLAG, C-ter, $Amp^R$ | This study |
| pEM15178 | pTrc99A-FF4-*fliI* (Δaa2-7, D341amber)-SAGASA-3×FLAG, C-ter, $Amp^R$ | This study |
| pEM15179 | pTrc99A-FF4-*fliI* (R177H, D341amber)-SAGASA-3×FLAG, C-ter, $Amp^R$ | This study |
| pEM15180 | pTrc99A-FF4-*fliI* (G183A, D341amber)-SAGASA-3×FLAG, C-ter, $Amp^R$) / pSUP (artificial amino-acid, $Cm^R$) | This study |
| pEM15181 | pTrc99A-FF4-*fliI* (E211D, D341amber)-SAGASA-3×FLAG, C-ter, $Amp^R$) / pSUP (artificial amino-acid, $Cm^R$) | This study |
| pEM15182 | pTrc99A-FF4-*fliI* (E211Q, D341amber)-SAGASA-3×FLAG, C-ter, $Amp^R$ | This study |
| pEM15183 | pTrc99A-FF4-*fliI* (E215Q, D341amber)-SAGASA-3×FLAG, C-ter, $Amp^R$ | This study |

**Table S2.** Oligonucleotides used in this study.

| Sequence | Source | Identifier |
|---|---|---|
| GGATTCTGTGGCGCAAAG | Lab Collection | 5'-fliFseq4-fw |
| TGGTTTTCAACGATGGCGTTAGC | Lab Collection | 3'-fliK95-YscP(138-353)_gBlock_rv |
| ATTACCGGCACGCTCGAC | Lab Collection | 3'-FliI-int900_rv |
| CCTGAAACCAAAGAGGTGGA | Lab Collection | 5'-FliI-int180_fw |
| TTACGCGAGAAGTTCCTGCG | Lab Collection | 5'_fliG-Cter-fw |
| CGCTCAATAATGCAGGTGG | Lab Collection | 3'-FliH-sequencing-rev |
| AACCGGCGCGTTAATCACGCCGCCGTTTAACC CGCTACAGAGGGTTTTCCCAGTCACGAC | Lab Collection | 5'_FliI-dR152-F181-KanSceI_fw |
| TCATGCCAAGCAGAACCGATTTACCAACGCCGG AACCGGCTGCTTCCGGCTCGTATGTTG | Lab Collection | 3'_FliI-dR152-F181-KanSceI_rv |
| CTACACGCGGGCGGACGTGATTGTCGTGGGAC TTATCGGCAGGGTTTTCCCAGTCACGAC | Lab Collection | 5'_fliI_KanSceI_dE211_fw |
| GACCGTCGGGGCCGAGAATATTTTCGATAAAAT CTTTAACTGCTTCCGGCTCGTATGTTG | Lab Collection | 3'_fliI_KanSceI_dE215_rv |
| TATTATTACGCCATCCGAGCGTATGCGTCGTTT GAGTCGTAGGGTTTTCCCAGTCACGAC | Lab Collection | FliN-KanSceI-Cter_FW |
| GCGGTGGGCTGAGAAACCGTGGCTTCTGTCTT CATCATTATGCTTCCGGCTCGTATGTTG | Lab Collection | FliN-KanSceI-Cter_RV |
| GTACTTCGCGGGTGGAGAG | This study | fliG-int-rev2 |
| AAGAGTTGTGTCGCCTGGCGGCGCCGGGAGTG CTCTGATGAGGGTTTTCCCAGTCACGAC | This study | FliI-N-ter-KanSce-fw |
| TTTTGGCTTCAAAGTTGTCGAGCGCGGTAAGCC AGCGGGTTGCTTCCGGCTCGTATGTTG | This study | FliI-N-ter-KanSce-rev |
| CCATTGCCTAGTCGGCACGCGCAATCCTCGAC GGG | This study | fliI D341amber fw |

| | | |
|---|---|---|
| GCGTGCCGACTAGGCAATGGGATCTTGTTGGT CGTCCCCCTC | This study | fliI D341amber rev |
| GGAATTGTGAGCGGATAACAATTTCACACAGGA AACAGCAATGACCACGCGCCTGACC | This study | GA_fliI_fw |
| AGCCAAGCTTGCATGCCTGCAGGTCGACTCTA GAGTCACTTGTCATCATCATCTTTATAA | This study | GA_fliI-3XFLAG_rev |
| ATGATGATGACAAGTGAGATCCTCTAGAGTCGA CCTGCAG | This study | GA_pTrc-fliI_fw |
| GGTCAGGCGCGTGGTCATTGCTGTTTCCTGTGT GAAATTGTTATCC | This study | GA_pTrc-fliI-3XFLAG_rev |
| GTGATTGGCAGCGGCGAGGATACCTATGTCTAA TGAATTGAGGGTTTTCCCAGTCACGAC | This study | DfliH-KanSceI-fw |
| CAGCGGGTCAGGCGCGTGGTCATCAGAGCACT CCCGGCGCTGCTTCCGGCTCGTATGTTG | This study | DfliH-KanSceI-rev |
| GTGATAAAGCAGGAGGGCGACGATCATGGCAC AACATGGCAGGGTTTTCCCAGTCACGAC | This study | DfliJ-KanSceI-fw |
| GTGATCAGTTGGGGCAGGGTGATCATTCGGGT TTCCTCATTGCTTCCGGCTCGTATGTTG | This study | DfliJ-KanSceI-rev |
| CCGACGGTGTGATAAAGCAGG | This study | DfliJ fw |
| CGGTCAGCATATGGGACTCTTTGCCGGTTCCG GCGTTGGTAAATCGG | This study | fliI-R177H onestep fw |
| GAGTCCCATATGCTGACCGCGCCCTACGGTTAA CAACGCGTTGATAGCGC | This study | fliI-R177H onestep rev |
| TTTGCCGCGTCCGGCGTTGGTAAATCGGTTCTG CTTGGCATGATGGCGCGC | This study | fliI-G183A onestep fw |
| CAACGCCGGACGCGGCAAAGAGTCCCATACGC TGACCGCGCCC | This study | fliI-G183A onestep rev |
| GGGACTTATCGGCGATCGTGGCCGCGAAGTTA AAGATTTTATCGAAAATATTCTCGGCCC | This study | fliI-E211D onestep fw |
| CACGATCGCCGATAAGTCCCACGACAATCACGT CCGCCCGC | This study | fliI-E211D onestep rev |
| GGGACTTATCGGCCAGCGTGGCCGCGAAGTTA AAGATTTTATCGAAAATATTCTCGGCCC | This study | fliI-E211Q onestep fw |
| CACGCTGGCCGATAAGTCCCACGACAATCACG TCCGCCCGC | This study | fliI-E211Q onestep rev |
| CGTGGCCGCCAGGTTAAAGATTTTATCGAAAAT ATTCTCGGCCCCGACGGTCG | This study | fliI-E215Q onestep fw |
| CTTTAACCTGGCGGCCACGTTCGCCGATAAGTC CCACGACAATCACGTCC | This study | fliI-E215Q onestep rev |
| GAAACAGCAATGTGGCTTACCGCGCTCGACAA CTTTGAAGCCAAAATGGCGTTATTG | This study | fliI-Daa2-7 one-step fw |
| CGGTAAGCCACATTGCTGTTTCCTGTGTGAAAT TGTTATCCGCTCACAATTCCACACATT | This study | fliI-Daa2-7 one-step rev |
| AAGAGTTGTGTCGCCTGGCGGCGCCGGGAGTG CTCTGATGTGGCTTACCGCGCTCGAC | This study | fliI-Daa2-7 replacement fw |
